# Tertiary lymphoid structures induced by CXCL13-producing CD4^+^ T cells increase tumor infiltrating CD8^+^ T cells and B cells in ovarian cancer

**DOI:** 10.1101/2021.12.01.470493

**Authors:** Masayo Ukita, Junzo Hamanishi, Hiroyuki Yoshitomi, Koji Yamanoi, Shiro Takamatsu, Akihiko Ueda, Haruka Suzuki, Yuko Hosoe, Yoko Furutake, Mana Taki, Kaoru Abiko, Ken Yamaguchi, Hidekatsu Nakai, Tsukasa Baba, Noriomi Matsumura, Akihiko Yoshizawa, Hideki Ueno, Masaki Mandai

## Abstract

**Background:** Tertiary lymphoid structures (TLSs) are transient ectopic lymphoid aggregates whose formation might be caused by chronic inflammation states, such as cancer. The presence of TLS is associated with a favorable prognosis in most solid malignancies. The recognition of the relevance of TLS to cancer has led to a growing interest in TLS as an immunomodulatory target to enhance tumor immunity, although how TLSs are induced in the tumor microenvironment (TME) and how they affect patient survival are not well understood.

**Methods:** TLS distribution in relation to tumor infiltrating lymphocytes (TILs) and related gene expression were investigated in high grade serous ovarian cancer (HGSC) specimens. CXCL13 expression, which is strongly associated with TLS, and its localization in immune cells, were examined. We explored the tumor microenvironment for CXCL13 secretion by adding various inflammatory cytokines *in vitro*. The induction of TLS by CXCL13 was examined in a mouse model of ovarian cancer.

**Results:** CXCL13 gene expression correlated with TLS formation and the infiltration of T cells and B cells, and was a favorable prognostic factor for HGSC patients. The coexistence of CD8^+^ T cells and B-cell lineages in the TME was associated with a better prognosis of HGSC and was closely related to the presence of TLSs. CXCL13 expression was predominantly coincident with CD4^+^ T cells in TLSs and CD8^+^ T cells in TILs, and shifted from CD4^+^ T cells to CD21^+^ follicular dendritic cells as the TLS matured. Although TGF-β was reported to stimulate CXCL13 production, our *in vitro* results revealed that CXCL13 secretion was promoted in CD4^+^ T cells under TGF-β + IL-2-restricted conditions and in CD8^+^ T cells under TGF-β + IL-12-rich conditions. In a mouse model of ovarian cancer, recombinant CXCL13 induced TLSs and enhanced survival by the infiltration of CD8^+^ T cells.

**Conclusions:** TLS formation was promoted by CXCL13-producing CD4^+^ T cells and TLSs facilitated the coordinated antitumor responses of cellular and humoral immunity in ovarian cancer.

## Background

It is generally considered that the generation and regulation of an efficient adaptive immune response to cancer occurs in secondary lymphoid organs (SLOs) such as the regional lymph nodes. Antitumor immune cells are educated to recognize tumor antigens, proliferate in regional lymph nodes away from the tumor site, and then migrate into the tumor microenvironment (TME) to exert antitumor activity (1). Clinical studies have shown that higher densities of T cell subsets within the TME are associated with improved patient survival in several cancers including ovarian cancer (2–5). We previously reported that the forced infiltration of CD8^+^ T cells into the TME by CCL19, a chemokine that attracts T cells, suppressed tumors in an ovarian cancer model (6). These results indicate that T cells have a critical role in the TME and that the TME might be a therapeutic target if effectively altered by immune-activating signals such as chemokines. To date, strategies to enhance the clinical efficacy of anti-tumor treatments have predominantly focused on the T cell component in the tumor, and the roles of other immune cell components have not been fully elucidated.

Recent studies have revealed an alternative immune response at the tumor site within SLO-like cellular aggregates called tertiary lymphoid structures (TLSs) (7). TLSs are transient ectopic lymphoid aggregates whose formation might be caused by chronic inflammation states, including autoimmune and infectious diseases, transplanted organ rejection, and cancer (7–9). The presence of TLS is associated with a favorable prognosis in most solid malignancies (10). Recently, it was reported that TLS-associated B cells synergized with T cells to contribute anti-tumor effects and that the presence of TLS and B cells in tumor sites enhanced the efficacy of immunotherapy (11–14). The recognition of the relevance of TLS to cancer has led to a growing interest in TLS as an immunomodulatory target to enhance tumor immunity, although how this can be induced therapeutically is not known.

The chemokines CXCL13, CCL19, and CCL21 are involved in lymphoid tissue-inducer (LTi) cell homing and lymph node development (8). Especially, CXCL13 was reported to be essential for the initial attraction of LTi cells and the formation of early lymph nodes (15). Although TLSs are thought to share the mechanisms of initial development with SLOs, TLS formation is distinct from the preprogrammed processes involved in SLOs and does not necessarily occur in all patients. The generation of TLS in inflamed tissues might be governed by specific inflammatory signals that have not been fully identified (7). In autoimmune diseases such as rheumatoid arthritis (RA), we showed that TGF-β and other proinflammatory cytokines enhanced CXCL13 production by CD4^+^ T cells, and that CXCL13-producing CD4^+^ T cells had an important role in the formation of TLS (16, 17). However, whether the same mechanism can be applied to the case of malignant tumors including ovarian cancer has not been investigated.

In this study, we assessed the relationship between tumor infiltrating T or B cells subsets and the presence of TLS, and evaluated the prognostic impact of TLS in ovarian cancer. We also investigated whether CXCL13 promoted TLS formation and improved the prognosis of patients with ovarian cancer. The role of CXCL13-producing CD4^+^ T cells in the generation of TLS was also investigated.

## Methods

### Human samples

Sixty-two and 35 high grade serous ovarian cancer (HGSC) patients who underwent primary surgery at Kyoto University Hospital from 1997 to 2015 and at Kindai University Hospital from 2009 to 2016, respectively, were enrolled. Their clinical characteristics are described in Supplemental Table S1. Patients who received chemotherapy or radiation therapy prior to surgery were excluded. Four other patients with typical TLSs who underwent initial surgery at the National Hospital Organization Kyoto Medical Center between 2017 and 2019 were also included in the study.

### Immunohistochemical analysis and evaluation

Immunohistochemical (IHC) staining was performed using formalin-fixed, paraffin-embedded (FFPE) specimens obtained from the above patients by the streptavidin-biotin-peroxidase method as previously described (4, 6). The samples were incubated with anti-CD8, anti-CD4, anti-CD20, anti-CD38, and anti-CD21 antibodies. The antibodies used are listed in Supplemental Table S2.

Two independent investigators trained in the pathology of ovarian cancer and blinded to the clinical data examined the H&E staining and IHC slides. TLSs were evaluated by H&E staining and CD20 positive cell aggregation as an indicator. Five sections at 400× magnification with the most abundant infiltration were manually counted and the mean count was calculated for CD8^+^ T cells, CD4^+^ T cells, and CD20^+^ B cells. These immune cells in TLSs were not counted as tumor infiltrating lymphocytes (TILs). If their count was above the median, we defined them as CD8-high, CD4-high, and CD20-high tumor, respectively. Tumor infiltrated CD38^+^ plasma cells were graded according to their intensity and fraction of positive cells as 0, 1, 2, or 3 (plasma cell score) according to previous reports (18, 19). Cases with scores of 0 and 1 were defined as plasma cell-low tumor, and cases with scores of 2 and 3 were defined as plasma cell-high tumors.

### Detection of CXCL13 mRNA by RNA ISH

We assessed CXCL13 by RNA in situ hybridization (ISH) (RNAscope^®^ 2.5 HD Reagent kit (RED), Advanced Cell Diagnostics, Hayward, CA, USA). FFPE tissue sections were deparaffinized in xylene and subsequently dehydrated in an ethanol series. Tissue sections were incubated in target retrieval reagent at 100°C for 15 minutes, and then treated with protease at 40°C for 30 minutes. Hybridization with Hs-CXCL13 (for human) or Mm-Cxcl13 (for mouse) probes at 40°C for 2 hours, and the amplifier and visualization (Fast RED) procedures were performed in accordance with the manufacturer’s instructions. For multiplex detection using FFPE, an RNAscope^®^ Fluorescent Multiplex Reagent kit v2 (Advanced Cell Diagnostics) was used. Double staining was performed for CXCL13 and CD8, CXCL13 and CD4, and CXCL13 and CD21(CR2). The target probes and reagents used were listed in Supplemental Table S2. CXCL13 was detected with Opal 690, and CD4, CD8, and CR2 with Opal 570. Fluorescence images were captured using a fluorescence microscope BZ-X800E (KEYENCE, Osaka, Japan), and the colocalization of CXCL13 with various immune cells was quantified using BZ-H4C/hybrid cell count software (KEYENCE).

### HGSC gene expression analysis

HGSC specimens obtained from 28 patients who underwent primary surgery at Kyoto University Hospital from 1997 to 2012 were prepared for gene expression microarray analysis (KOV). The data were previously deposited in the Gene Expression Omnibus (Accession Numbers: GSE 39204 and GSE 55512).

The gene expression profile of the TCGA-OV RNA sequencing dataset (n=217) from The Cancer Genome Atlas (TCGA) Data Portal (http://cancergenome.nih.gov, illuminahiseq_rnaseqv2_Level_3_RSEM_genes_normalized_data files obtained and merged on 19, Oct, 2015) was used for survival analysis and correlation testing among CXCL13, TGF-β1, PDCD1, and CD274.

### Gene expression and infiltrating immune cells analysis

Raw gene expression microarray data using Affymetrix HT_HG-U133A from TCGA ovarian serous cystadenocarcinoma samples were obtained from the GDC legacy archive (https://portal.gdc.cancer.gov/legacy-archive/) in the form of CEL files (n=522). Gene expression values were calculated by normalization using the RMA method with R package “affy” (http://www.R-project.org). Subsequently, the relative abundance of 22 immune cell types for each sample was estimated using CIBERSORT (http://cibersort.stanford.edu/).

### T cell receptor (TCR) and B cell receptor (BCR) repertoire analysis

TLSs and tumor areas were identified by microscopy and macrodissected independently from FFPE sections (Supplemental Figure S1C). RNA was extracted using NucleoSpin^®^ total RNA FFPE (Takara Bio, Shiga, Japan) according to the manufacturer’s instructions. Sequencing of the TCRα (TRA) and BCR IgG heavy chain loci were performed at Repertoire Genesis Incorporation (Osaka, Japan) using an unbiased amplification method with MiSeq (Illumina, San Diego, CA, USA). Data processing, assignment, and aggregation were performed using a repertoire analysis software program, Repertoire Genesis (RG), provided by Repertoire Genesis Incorporation. RG assigns TRV and TRJ alleles to queries and then generates CDR3 sequences, finally aggregating their combination patterns.

### Induction assay of CXCL13 in CD4^+^ and CD8^+^ T cells

PBMCs from healthy donors were collected using Lymphocyte Separation Solution 1.077 (Nacalai Tesque, Kyoto, Japan). Blood CD8^+^ T cells were isolated with CD8 MicroBeads, human (Miltenyi Biotec, Bergisch Gladbach, NRW, Germany). Blood CD4^+^ T cells were purified with a Naïve CD4^+^ T cell isolation kit II, human (Miltenyi Biotec) through a magnetic column.

Human T cells were differentiated for 6–7 days in a humidified 5% CO_2_ incubator at 37°C with IMDM (Thermo Fisher, Waltham, MA, USA) supplemented with 10% fetal bovine serum, 100 units/ml penicillin and streptomycin under stimulation with 5 µg/ml plate-bound CD3 monoclonal antibody (clone: OKT3, Thermo Fisher) and 10 µg/ml CD28 monoclonal antibody (clone: CD28.2, Thermo Fisher) in the presence of 10 ng/ml TGF-β1 (Cell Signaling Technology, Danvers, MA, USA) unless otherwise described. Human T cells were also cultured under conditions where 5 µg/ml neutralizing anti-IL-2 antibody (R&D Systems, Minneapolis, MN, USA) or 10 ng/ml IL-12 (PeproTech, Montreal, Quebec, Canada) was added.

Human T cells were cultured under CD3/CD28 stimulation using conditioned medium from the human serous ovarian cancer cell lines: OVCA420, OVCA433, and DK-09. OVCA420 and OVCA433 were kindly provided by Dr. Susan K. Murphy of Duke University. DK-09 is a cell line that we established from the ascites of a patient with recurrent ovarian cancer (see Supplemental Methods). To block the TGF-β signals, a TGF-β signal inhibitor, SB431542 (Stemgent, Beltsville, MD, USA) was added at 0.5 and 5 µM.

### Flow cytometry

For intracellular staining, cells were cultured for 4 hours with 4 µM monensin (Sigma-Aldrich, Saint Louis, MO, USA), fixed, and stained with eBioscience™ Intracellular Fixation & Permeabilization Buffer Set (Thermo Fisher) and antibodies for intracellular molecules. Fixable Viability Dye eFluor 506 (Thermo Fisher) was used to exclude dead cells. To detect CXCL13, CXCR5 and PD-1, the antibodies listed in supplemental table S2 were used. Data were acquired using MACS Quant Analyzer 10 (Miltenyi Biotec) and were analyzed with FlowJo 10.0 (FlowJo LLC, Ashland, OR, USA).

### ELISA

The concentrations of CXCL13 and TGF-β1 in the supernatant were measured with the respective kits listed in Supplemental Table S2.

### Cell lines and tumor models

The OV2944-HM-1(HM-1) mouse ovarian cancer cell line was purchased from RIKEN BioResource Center (Ibaraki, Japan) and cultured as described (20). Throughout the study, we used HM-1 cell lines passaged fewer than 20 times, and regularly tested for mycoplasma contamination. Female B6C3F1 (C57BL6 × C3/He F1) mice and nude mice (BALB/C-nu: CAnN.Cg-*Foxn1^nu^*/Crl) were purchased from Charles River Japan (Yokohama, Japan), and were maintained under specific pathogen-free conditions.

A total of 1 × 10^6^ HM-1 cells were inoculated intraperitoneally into B6C3F1 mice. Mouse recombinant CXCL13 (R&D Systems) treatment was initiated one day after the tumor inoculation and administered intraperitoneally at 1 µg/mouse every other day for five times. Control mice received PBS intraperitoneally. Then, 10–12 days after inoculating the tumor, mice were euthanized with carbon dioxide and the formation of TLS in omental tumors was analyzed.

A total of 2.5 × 10^5^ HM-1 cells were inoculated intraperitoneally into B6C3F1 and nude mice. Similarly, rCXCL13 (1 µg/mouse) and anti-PD-L1 antibody (200 µg/mouse) were intraperitoneally administered 5 times every other day starting from day 1 and day 3, respectively, after tumor implantation. Anti-PD-L1 antibody (clone 10F.9G2, Bio X Cell, Lebanon, NH, USA) and Rat IgG antibody (clone LFT-2, Bio X Cell) were used as negative controls.

### IHC analysis of mouse tumors

Mouse tumor cryosections (6-µm-thick) were stained with anti-CD4, anti-CD8 (clone YTS169.4), anti-CD19, and anti-Ki-67 antibodies as previously described (21). The antibodies are listed in Supplemental Table S2. Mouse FFPE specimens were stained with anti-CD8 antibody (clone EPR20305).

### Statistics

Results are shown as the mean ± SEM from at least three independent experiments unless otherwise stated. A *P* value of less than 0.05 was considered statistically significant. Significance was calculated using the 2-tailed Student’s *t*-test, and correlation between groups was determined by Spearman’s correlation test. The log-rank test was used for overall survival analysis unless otherwise described. All statistical analyses were performed using GraphPad Prism 7 (GraphPad software, San Diego, CA, USA). The Jonckheere-Terpstra test was performed using the “clinfun” packages in R.

## Results

### TLS is associated with intratumor infiltration by CD8^+^ T cells and B-cell lineages and is closely related to a favorable prognosis

The prognostic significance of the respective infiltration of T cell and B cell subsets was investigated by the IHC analysis of initial surgical specimens of HGSC (n=97) (Supplemental Table S1). Patients with a higher infiltration of CD8^+^ T cells, CD20^+^ B cells, and CD38^+^ plasma cells had significantly prolonged progression free survival than those with low numbers of infiltrating cells (*P*<0.05, each) (Figure 1A). In addition, there was a significant correlation between the infiltrated number of CD8^+^ T cells and B-cell lineage cells (Figure 1B), and patients with a higher number of infiltrated CD8^+^ T cells and B-cell lineages had the best prognosis (Figure 1C). The intratumoral infiltration of CD8^+^ T cells or B-cell lineages alone did not contribute to the improved prognosis.

**Figure 1.**
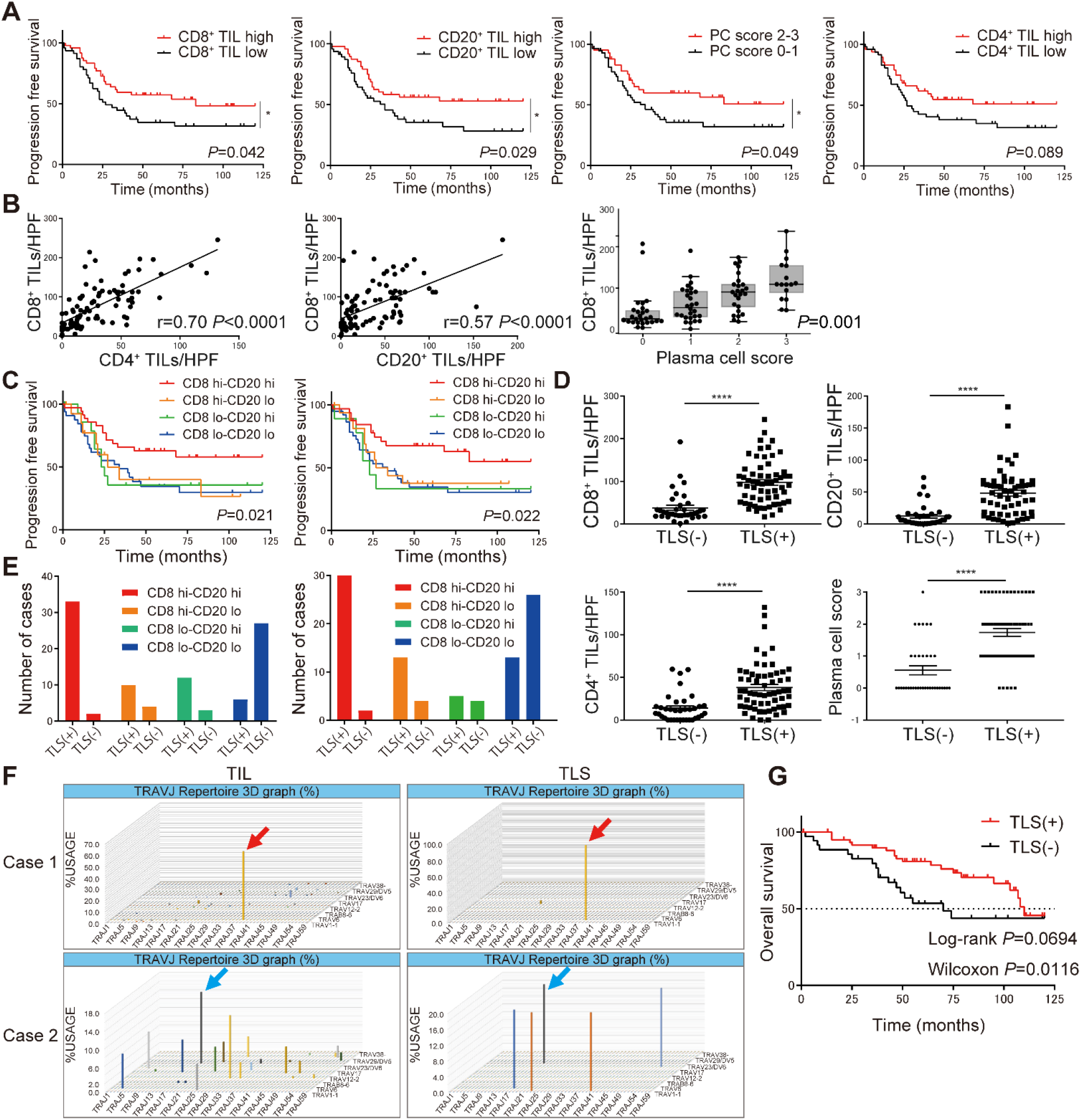
TLS is associated with intratumor infiltration by CD8^+^ T cells and B-cell lineages and is closely related to a favorable prognosis. **(A)** Progression free survival of the cohort stratified by CD8^+^ T cells, CD20^+^ B cells, plasma cells (PC), and CD4^+^ T cells (n=97, each). **(B)** Association between infiltrated numbers of CD4^+^ T cells and CD8^+^ T cells, and CD8^+^ T cells and B cell lineages in tumors (n=97). Correlations were determined by Pearson’s correlation test and Jonckheere-Terpstra trend tests. **(C)** Progression free survival of patients based on the tumor-infiltrating CD8^+^ T cells and B lineage cells (CD8 hi-CD20 hi: n=35, CD8 hi-CD20 lo: n=14, CD8 lo-CD20 hi: n=15, CD8 lo-CD20 lo: n=33, total n=97) (CD8 hi-PC hi: n=32, CD8 hi-PC lo: n=17, CD8 lo-PC hi: n=9, CD8 lo-PC lo: n=39, total n=97). **(D)** The number of tumor-infiltrating CD8^+^ T cells, CD20^+^ B cells, plasma cells, and CD4^+^ T cells according to TLS presence (TLS− n=36, TLS+ n=61). *P* values were determined by Mann-Whitney *U-*test. **(E)** Distribution of TLS in relation to the infiltration pattern of immune cells in tumors. Tumors were considered high (hi) for CD8^+^ T cells, CD20^+^ B cells and CD4^+^ T cells if their score was above the median. Tumors were divided into two groups for PC by plasma cell score (0–1 n=56, 2–3 n=41), with 0–1 defined as PC low (lo) and 2–3 as PC hi in (A), (C), and (E). **(F)** TCR repertoire analysis separating TLS and TIL regions. The horizontal axis shows the J gene, the depth shows the V gene, and the vertical axis shows the frequency of usage. Clones indicated by arrows of the same color confirm the same amino acid sequence of CDR3. **(G)** Overall survival of patients with HGSC by the presence of TLS (TLS− n=36, TLS+ n=61, total n=97). Analyses were performed with Kaplan-Meier estimates and log-rank tests in (A) (C), and Wilcoxon tests (G).

H&E and IHC staining evaluation of TLSs in the same samples (n=97) revealed that 61 patients (62.9%) had TLSs (Supplemental Table S1, Supplemental Figure S1, A and B). In the cases with TLS, the number of infiltrating CD8^+^ T cells, CD4^+^ T cells, CD20^+^ B cells, or CD38^+^ plasma cells (plasma cell score) was significantly higher than in those without TLS (*P*<0.0001, each) (Figure 1D). Focusing on the pattern of tumor infiltrating lymphocytes and the presence of TLS, TLS were found in 94% of cases with a high infiltration of CD8^+^ T cells and B-cell lineages (CD8 high-CD20 high: n=35, CD8 high-plasma cell high: n=32) (Figure 1E). The close relationship between TLS and the distribution of infiltrating CD8^+^ T cell and B-cell lineages suggests that cellular and humoral immunity interact via the TLS in ovarian cancer.

Next, we performed TCR and BCR repertoire analysis using tumor sections from HGSC patients (n=3), separating TLS and TIL regions by macrodissection (Supplemental Figure S1C). In two of three cases, TCR repertoire analysis showed many clones distributed in the TIL, whereas oligoclonal amplification was observed in TLS. Furthermore, the clone with the highest amplification from TIL was consistent with the clone observed in TLS (Figure 1F), suggesting that antigen-specific T cells that proliferated in TLS might also infiltrate into the tumor as TIL.

Patients with TLS, which is closely associated with intratumoral infiltration of CD8^+^ T cells and B-cell lineages and may mediate cellular and humoral immunity, had a significantly better prognosis than patients without TLS (Wilcoxon test *P*=0.0016, median overall survival [110 months vs 70 months]) (Figure 1G).

### CXCL13 gene expression in tumors correlates with TLS formation, lymphocyte infiltration, and a favorable prognosis for ovarian cancer

The expression of CXCL13 analyzed by RNA ISH demonstrated CXCL13 was highly expressed in TLS (Figure 2A). In addition, there were cases in which immune cells in the tumor stroma also expressed high levels of CXCL13 (Supplemental Figure S2A). From the IHC of initial surgical specimens with our original microarray data (KOV: GSE39204/55512, n=28), the ratio of cases with TLS was significantly higher in those with high CXCL13 gene expression than in those with low CXCL13 gene expression in tumor specimens(*P*=0.046) (Figure 2B). Additionally, CXCL13 gene expression was significantly correlated with the numbers of several types of TILs such as CD4^+^ and CD8^+^ T cells, CD20^+^ B cells and CD38^+^ plasma cells (*P*<0.001, each) (Figure 2C, Supplemental Figure S2B).

**Figure 2.**
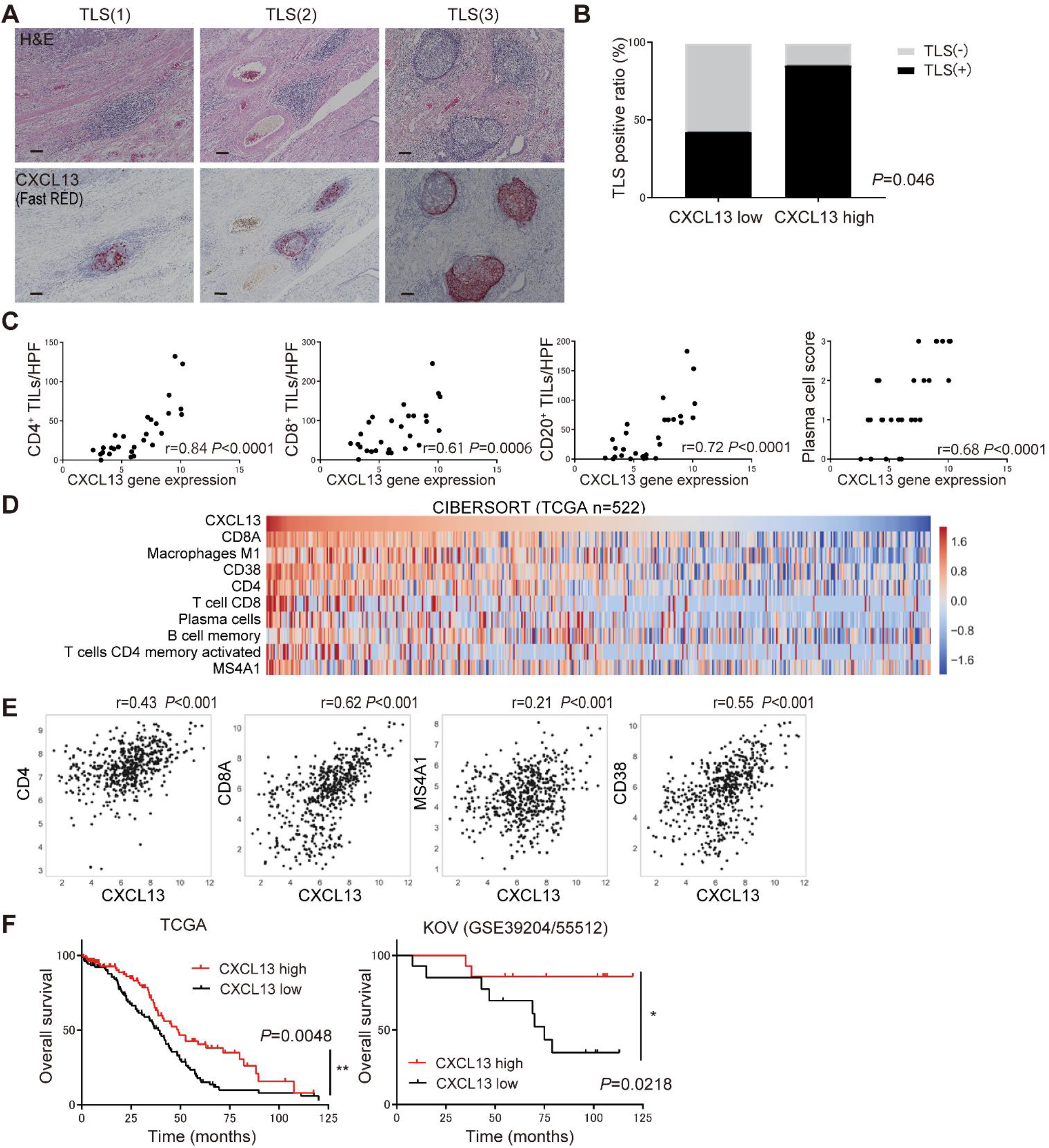
CXCL13 gene expression in tumors correlates with TLS formation, lymphocyte infiltration, and a favorable prognosis for ovarian cancer. **(A)** Representative TLS in tissues stained by H&E and CXCL13 (Fast RED) by RNA ISH. Scale bars indicate 100 µm. **(B)** TLS presence ratio based on CXCL13 gene expression. Analysis by Fisher’s exact test in 28 cases with microarray data. **(C)** Characterization of the immune infiltrate in tumors according to CXCL13 gene expression (n=28). Correlation was determined by Spearman’s correlation test. **(D) (E)** The distribution of infiltrating immune cells into the tumor site and CXCL13 gene expression using CIBERSORT (n=522). Correlation was determined by Spearman’s correlation test. **(F)** Overall survival of patients with HGSC by CXCL13 gene expression (TCGA n=217, KOV n=28). Patients with CXCL13 high defined if CXCL13 gene expression was above the median. Analyses were performed with Kaplan-Meier estimates, log-rank tests and Wilcoxon tests. The level of significance was set as \**P*<0.05, \*\**P*<0.01, and \*\*\*\**P*<0.0001.

To validate these data, we applied CIBERSORT to examine the distribution of infiltrating immune cells into tumor sites and CXCL13 gene expression using the RNA sequence data of ovarian cancer cases registered in TCGA. The infiltration of CD8^+^ T cells had the strongest correlation with CXCL13 gene expression, and that of CD4^+^ T cells, CD20^+^ B cells, CD38^+^ plasma cells and M1-macropahges also showed a strong correlation with CXCL13 gene expression, while that of M2-macrophages and mature dendritic cells were negatively correlated, and that of natural killer cells and regulatory T (Treg) cells were not correlated with CXCL13 gene expression (Figure 2, D and E, Supplemental Figure S2, C and D). These results suggest that CXCL13 gene expression strongly correlated with the formation of TLS and the number of tumor-infiltrating T cells and B-cell lineages.

Next, we examined the impact of CXCL13 on the prognosis of HGSC and found that patients with high CXCL13 gene expression had a significantly better prognosis in the KOV data and TCGA data (*P*<0.05, each) (Figure 2F).

### CXCL13 produced by CD4^+^ T cells is critical for TLS initiation

To identify cells producing CXCL13 involved in TLS formation, we performed RNA ISH double staining (CXCL13 and CD4^+^ T cells, CXCL13 and CD8^+^ T cells) using HGSC tumor specimens. In the TLS region, CXCL13 was highly coexpressed with CD4^+^ T cells, whereas CD8^+^ T cells predominantly expressed CXCL13 in the tumor and stromal regions (Figure 3, A and B).

**Figure 3.**
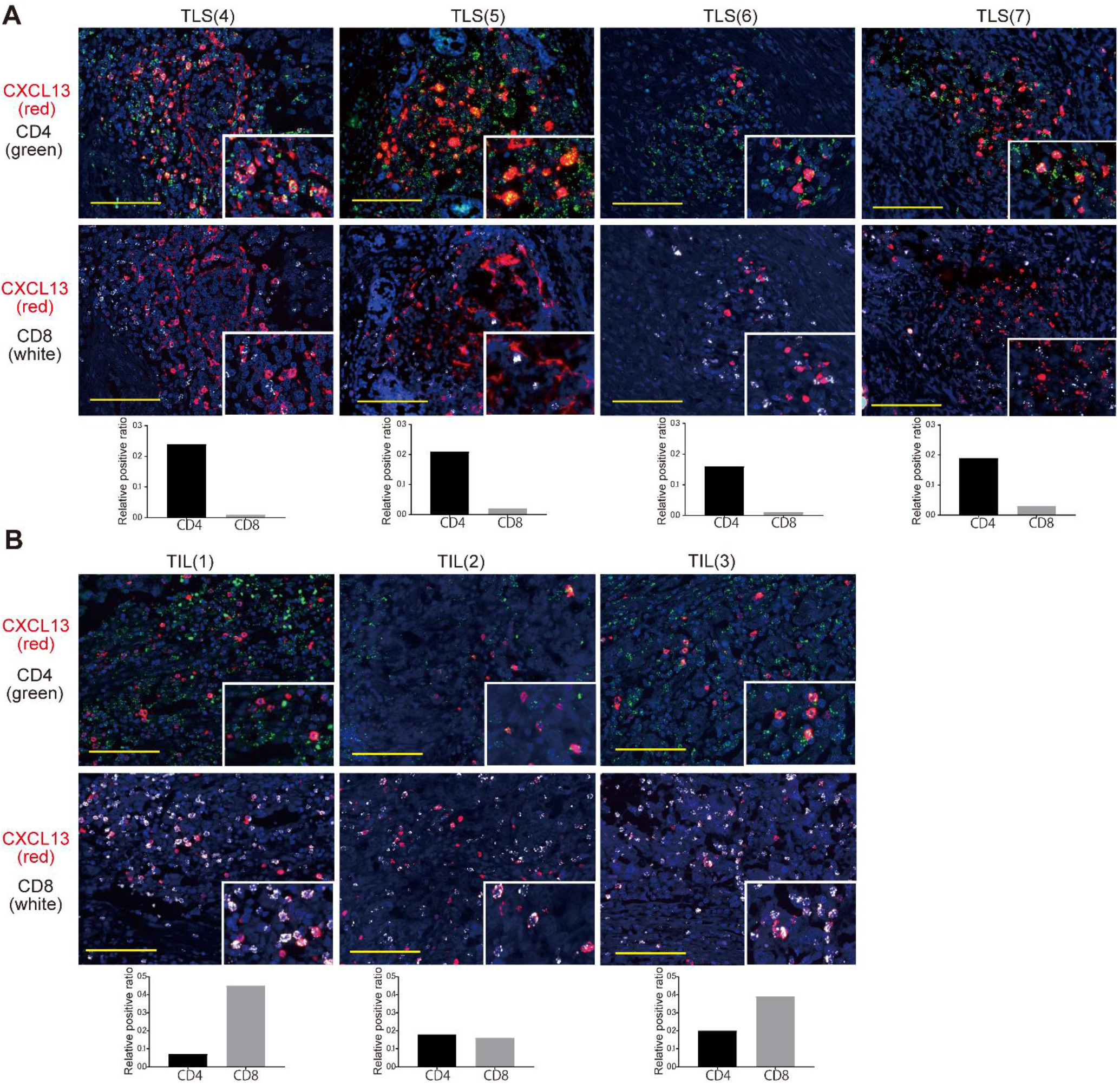
CXCL13 is mainly produced by CD4^+^ T cells in TLS. **(A)** Fluorescent double staining of CXCL13 (red) and CD4 (green), and CXCL13 (red) and CD8 (white) by RNA ISH in TLS. Images of four representative TLS are shown. **(B)** Fluorescent double staining of CXCL13 (red) and CD4 (green), and CXCL13 (red) and CD8 (white) by RNA ISH in TIL. The upper and lower pictures are representative TIL images from the same patient. Nuclei are stained with DAPI (blue). Scale bars indicate 100 µm. Co-localization of CXCL13 with CD4 or CD8 is shown in the bar graph as the relative positive ratio quantified using BZ-H4C/hybrid cell count software.

HGSC tissues contain two types of TLSs: early TLS, in which lymphocytes aggregate diffusely and CD21^+^ cells are scarce, and follicle-formed TLS, which has the follicular morphology of SLO and where CD21^+^ follicular dendritic cells (FDCs) are distributed in a reticular pattern (Figure 4, A and B). Therefore, we performed the double staining of CXCL13 and CD8^+^ T cells, CXCL13 and CD4^+^ T cells, and CXCL13 and CD21^+^ FDCs by RNA ISH for representative early TLS and follicle-formed TLS. CXCL13 expression was highly consistent with CD4^+^ cells in early TLSs, whereas few CD4^+^ T cells expressing CXCL13 were observed in follicle-formed TLSs and CXCL13 expression was highly consistent with spindle-shaped CD21^+^ FDCs (Figure 4C). These results indicate that CXCL13-producing CD4^+^ T cells are closely related to the early stage of TLS formation.

**Figure 4.**
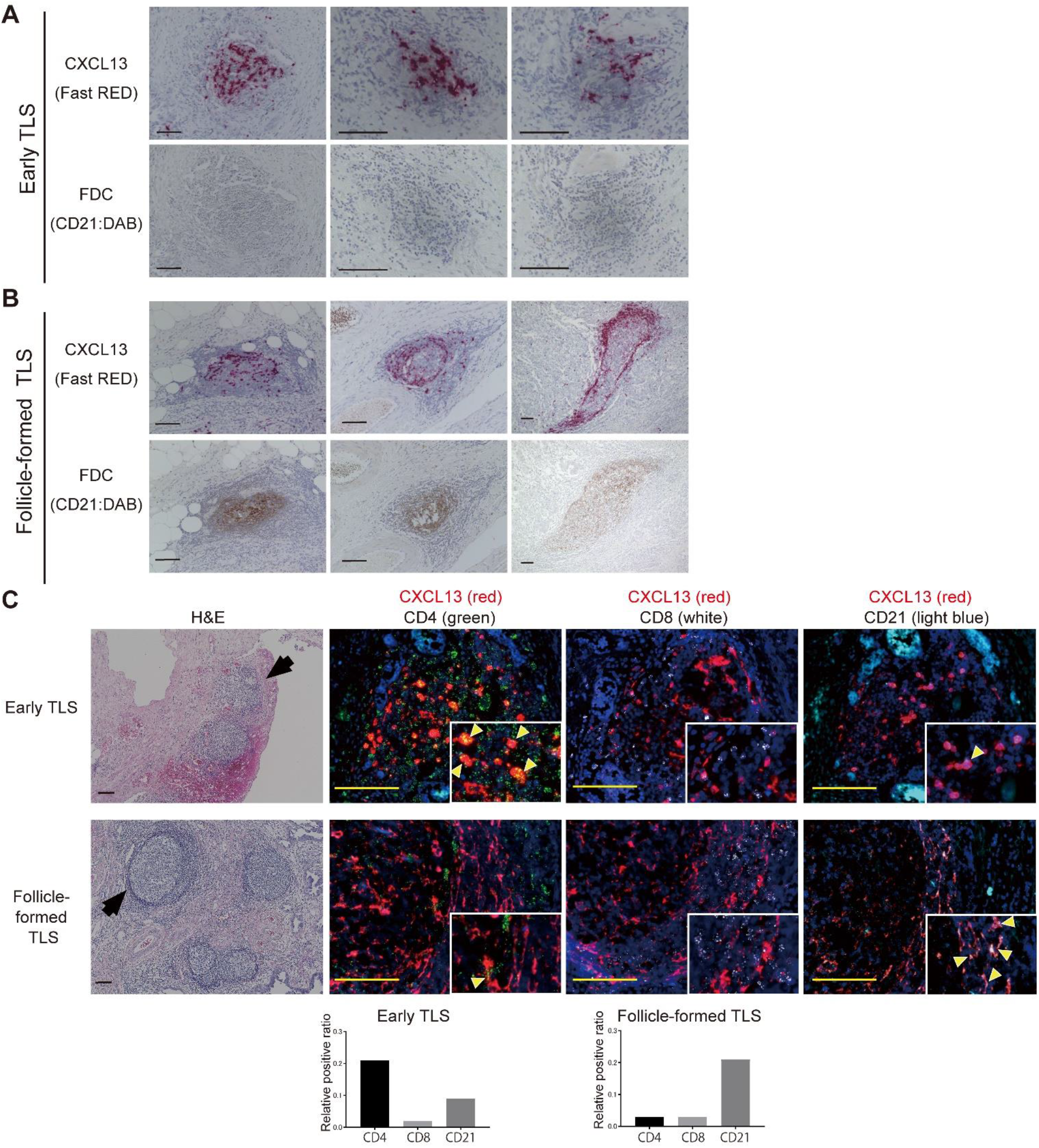
The source of CXCL13 production in TLS shifts to CD21^+^ FDC with the maturation of TLS. **(A)** Representative images of early TLS. **(B)** Representative images of follicle-formed TLS. Upper panels show CXCL13 (RNA ISH, Fast RED) and lower panels show FDC (CD21 IHC, DAB). **(C)** Fluorescence double staining of CXCL13 (red) and CD4 (green), CXCL13 (red) and CD8 (white), and CXCL13 (red) and CD21 (light blue) in representative early TLS and follicle-formed TLS. Nuclei are stained with DAPI (blue). Scale bar indicates 100 µm. Co-localization of CXCL13 with CD4, CD8, or CD21 is shown in the bar graph as the relative positive ratio quantified using BZ-H4C/hybrid cell count software.

### TGF-β promotes the production of CXCL13

To investigate which factors promote CXCL13 secretion from CD4^+^ T cells and CD8^+^ T cells, we analyzed two sets of gene expression data from ovarian cancer tissues. Using the TCGA RNA sequence data and our KOV microarray data, we found a significant correlation between CXCL13 and TGF-β1 gene expression (*P*<0.05) (Figure 5A). The analysis of naïve CD4^+^ T cells and CD8^+^ T cells isolated from the peripheral blood cells of a healthy donor and cultured with TGF-β showed that CD4^+^ T cells predominantly secreted CXCL13 compared with CD8^+^ T cells. A TGF-β signal inhibitor (SB431542) suppressed the secretion of CXCL13 in a concentration-dependent manner (Figure 5B).

**Figure 5.**
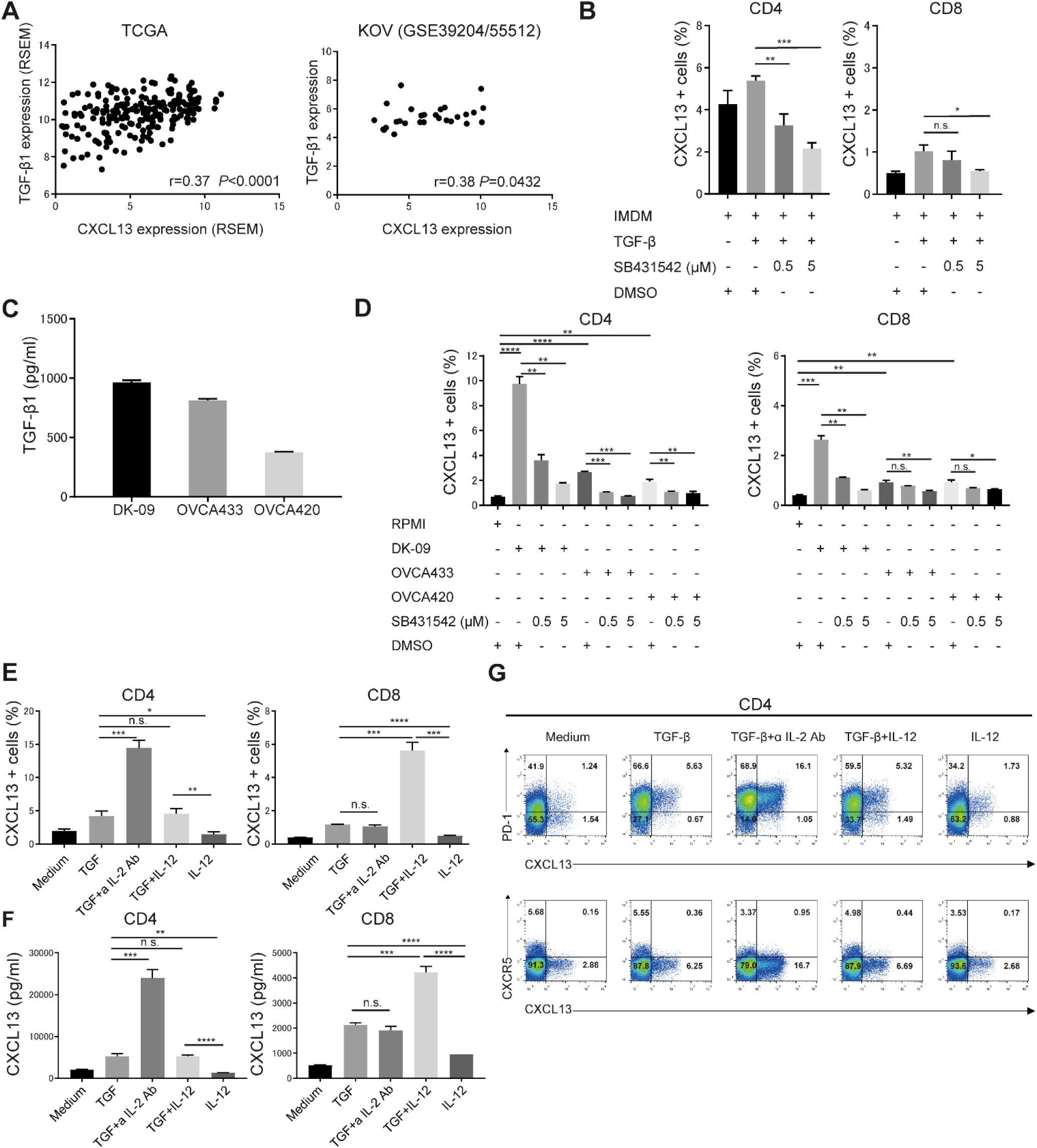
TGF-β promotes the production of CXCL13. **(A)** Correlation between CXCL13 and TGF-β1 expression in TCGA (n=217) and KOV (n=28). Correlation was determined by Spearman’s correlation test. **(B)** Human naïve CD4^+^ and CD8^+^ T cells from a healthy donor were differentiated by TCR stimulation and TGF-β1 in the presence or absence of a TGF signal inhibitor, SB431542. The proportion of CXCL13^+^ cells was determined by flow cytometry. Data are shown as the mean ± SEM of four samples. Statistical significance was determined by two-tailed Student’s *t*-test, \**P*<0.05, \*\**P*<0.01, \*\*\**P*<0.001, n.s.: not significant. **(C)** The concentration of TGF-β1 in conditioned medium obtained from three human ovarian cancer cell lines was measured by ELISA. Data are shown as the mean ± SEM of three samples. **(D)** Human naïve CD4^+^ and CD8^+^ T cells from a healthy donor were differentiated with TCR stimulation and conditioned medium obtained from three human ovarian cancer cell lines in the presence or absence of a TGF signal inhibitor, SB431542. The proportion of CXCL13^+^ cells was determined by flow cytometry. Data are shown as the mean ± SEM of triplicates. **(E) (F) (G)** Human naïve CD4^+^ and CD8^+^ T cells from a healthy donor were differentiated with TCR stimulation and the indicated cytokines. The proportion of CXCL13^+^ cells was determined by flow cytometry **(E)**. The concentration of CXCL13 in the culture supernatant was measured by ELISA **(F)**. Data are shown as the mean ± SEM of four samples in CD4 and three samples in CD8. Statistical significance was determined by two-tailed Student’s *t*-test, \**P*<0.05, \*\**P*< 0.01, \*\*\**P* < 0.001, \*\*\*\**P*<0.0001, n.s.: not significant. Representative dot plots of PD-1 (upper row), CXCR5 (lower row), and intracellular CXCL13 in healthy human naïve CD4^+^ T cells are shown **(G)**. a IL-2 Ab indicates anti IL-2 antibody.

We previously reported that the TGF-β signaling pathway was activated in ovarian cancer (22), and we detected high concentrations of TGF-β1 in the conditioned medium of three different human ovarian cancer cell lines (Figure 5C). Using these conditioned media, we examined their effects on CXCL13 production in naïve CD4^+^ and CD8^+^ T cells. In CD4^+^ T cells, CXCL13 secretion was promoted in the order of TGF-β1 concentration, and was suppressed in a concentration-dependent manner when incubated with the TGF-β signal inhibitor (SB431542) (Figure 5D). However, TGF-β-mediated CXCL13 secretion in CD8^+^ T cells was limited.

Next, we conducted similar experiments by adding various cytokines to TGF-β to reproduce the TME. We found that CXCL13 secretion was enhanced in CD4^+^ T cells under IL-2-restricted conditions and in CD8^+^ T cells under IL-12-enriched conditions (Figure 5E). The CXCL13 concentration in the culture supernatant showed a similar trend (Figure 5F). CXCL13-producing CD4^+^ cells had a PD-1 positive, CXCR5 negative phenotype (Figure 5G). These results suggest that the phase and TME in which CD4^+^ and CD8^+^ T cells produce CXCL13 are different, and thus CD4^+^ T cells may produce CXCL13 in TLSs and CD8^+^ T cells in TILs (Figure 3, A and B).

### Mouse recombinant CXCL13 induces TLS in tumors and prolongs survival

The effect of CXCL13 on TLS formation in tumors was analyzed using a mouse ovarian cancer model. A mouse ovarian cancer cell line HM-1 was intraperitoneally inoculated to B6C3F1 immunocompetent mice, and mouse recombinant(r) CXCL13 was intraperitoneally administered 5 times every other day starting from day 1, and TLS formed in the omental tumor were evaluated on days 10–12. The area of TLSs per tumor area was significantly increased in the rCXCL13-treated group compared with the control group (Figure 6A). IHC revealed that mouse TLS, similar to human TLS, consisted mainly of CD19^+^ B cells, and that CD8^+^ T cells and CD4^+^ T cells were present in and around TLSs. Furthermore, TLSs contained many Ki-67 positive immune cells indicating these structures were immunologically activated (Supplemental Figure S3A). Furthermore, CXCL13 was highly expressed and corresponding with TLSs (Figure 6B). The administration of rCXCL13 markedly increased the infiltration of CD8^+^ T cells around the TLSs (Figure 6C).

**Figure 6.**
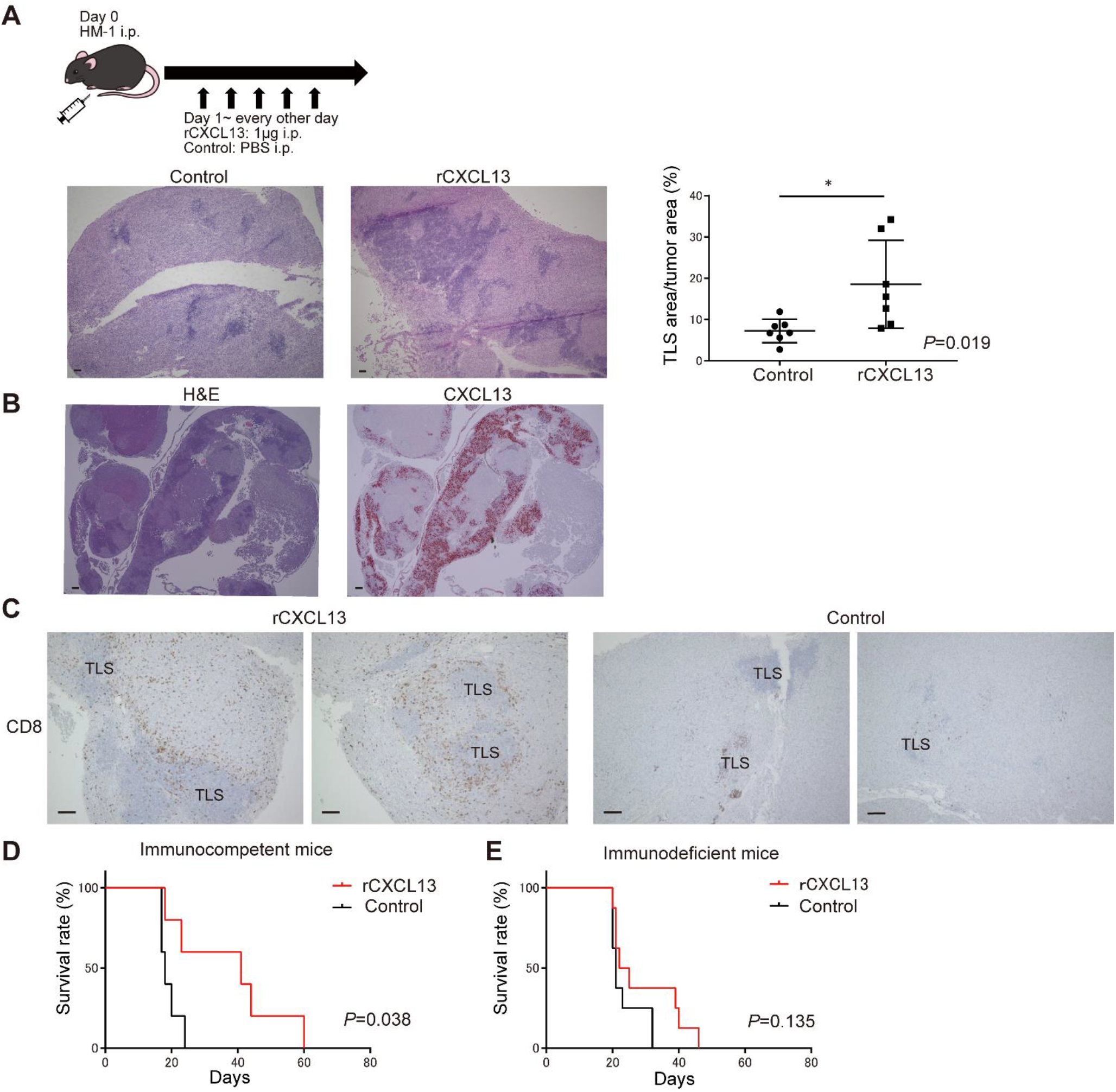
Mouse recombinant CXCL13 induces TLS in tumors and prolongs survival. **(A)** Mouse rCXCL13 was administered intraperitoneally to induce TLS in a mouse ovarian cancer model. Representative H&E images of TLS formed in an omental tumor. The area of TLS per tumor area was compared between the control group and the rCXCL13 treated group (n=7, each). Statistical significance was determined by two-tailed Student’s *t*-test, \**P*<0.05. **(B)** TLS induced by mouse rCXCL13 (H&E) and expression of mouse CXCL13 corresponding to TLS (RNA ISH, Fast RED). **(C)** CD8^+^ T cell IHC images (DAB) in the rCXCL13 treated group and control group. Scale bars indicate 100 µm. **(D) (E)** The effect of rCXCL13 administration on the survival of tumor-bearing mice was compared between immunocompetent mice (D) and immunodeficient mice (E). Analyses were performed using Kaplan-Meier estimates and log-rank tests.

Next, we observed the effect of rCXCL13 administration on the survival of mice. In immunocompetent mice (B6C3F1), the survival time was significantly prolonged in the rCXCL13-treated group compared with the control group (Figure 6D). Because there was a correlation between CXCL13 and PD-1/PD-L1 gene expression (Supplemental Figure S3B), we hypothesized that CXCL13 had an adjuvant effect in HM-1 ovarian cancer models that were originally refractory to anti-PD-1/PD-L1 antibody therapy. However, no significant prognostic improvement was observed (Supplemental Figure S3C).

Furthermore, the administration of rCXCL13 to immunocompromised mice (nude mice) did not improve survival (Figure 6E). Consistent with the apparent increase in CD8^+^ T cells around the TLSs in immunocompetent mice, CXCL13 contributed to their improved survival via immunity. These results indicate that CXCL13 induced TLSs and TILs, indicating CXCL13 and TLS might be new therapeutic targets for ovarian cancer.

## Discussion

We showed that the presence of TLS was associated with an increased intratumor infiltration of CD8^+^ T cells and B-cell lineages, and that CD8^+^ T cells and B-cell lineages might cooperate to improve the prognosis of ovarian cancer. It was reported that tumor infiltrating B cells contributed to tumor growth and progression through the production of cytokines, such as IL-10, that inhibit antitumor immunity although the functional role of B cells in cancer is poorly understood (23). Recently, there have been increasing numbers of reports that the presence of B cells, especially those associated with TLSs, may improve cancer outcomes (10, 12–14). Tumors containing CD8^+^ T cells and B-cell lineages were associated with improved prognosis in melanoma, sarcoma, and ovarian cancer (12, 13, 18). In this study, we found a strong correlation between the infiltrated numbers of CD8^+^ T cells and B-cell lineages, and confirmed the presence of TLS in 94% of patients with tumors containing high numbers of infiltrated T and B cells. These results suggest that B-cell lineages are an essential component of the CD8^+^ T cell cytotoxic reaction, and that cellular and humoral immune interactions are mediated by TLSs.

Whether tumor-associated TLSs are formed in response to a series of chronic inflammation or whether they are induced as a tumor antigen-specific immune response has not been fully established. De Chaisemartin et al. demonstrated that TLSs provided the specialized vasculature and chemoattractants necessary for T cell infiltration into non-small cell lung cancer (NSCLC) (24). The TCR repertoire analysis of NSCLC showed that the expansion of T cell clones in the tumor bed and peripheral blood correlated with the density of tumor associated TLSs (25). In this study, TCR repertoire analysis of the TLSs and tumor regions revealed that oligoclonal expansion occurred in TLS and that the same clone was highly amplified in the tumor. These data suggest that the recognition of antigens occurs in TLS and that the effector T cell clones amplified by the TLS infiltrate into tumors, both of which might provide evidence that TLS promotes immune responses in TME.

High CXCL13 gene expression was associated with disease activity and pathogenicity in autoimmune diseases such as RA (26–28), and with patients’ prognosis in several types of cancer (7, 29, 30), implying the strong involvement of CXCL13-dependent TLS formation. In line with previous reports, CXCL13 gene expression is a prognostic factor for ovarian cancer and is strongly associated with the formation of TLS. The presence of TLS also improved the long-term prognosis of ovarian cancer.

We previously reported that CXCL13-producing PD-1 high CXCR5 negative CD4^+^ T cells have an important role in the function of TLS in RA (16, 17). Although FDCs are the main source of CXCL13 in SLOs (31), the origin of CXCL13 in tumor associated TLS depends on the type of cancer. CXCL13 was secreted by PD-1^high^ CD8^+^ T cells in lung cancer (32), by CD103^+^ CD8^+^ cells in ovarian cancer (33), and by CXCR5^-^ PD-1^high^ CD4^+^ follicular helper like T cells in breast cancer (34). In this study, CXCL13 was predominantly expressed on CD4^+^ T cells in TLSs and on CD8^+^ and CD4^+^ T cells in TILs, and that the expression of CXCL13 in TLSs shifted from CD4^+^ T cells to CD21^+^ FDCs. Sequential stages of the development in tumor-associated TLS were observed in lung cancer. Siliņa et al. defined three types of TLSs: early TLS (E-TLS) without FDCs or germinal centers (GC), primary follicle like TLS (PFL-TLS) with an FDC network and lacking GC, and secondary follicle like TLS (SFL-TLS) with an FDC network and GC formation (35). Our data suggest that CXCL13-producing CD4^+^ T cells are an important primary producer of CXCL13 in the early stages of TLS when FDCs are not present. FDCs emerge from ubiquitous perivascular mesenchymal cells expressing platelet-derived growth factor receptor β (36). CXCL13 itself directly induces lymphotoxin (LT) production by naïve B cells, and this CXCL13/LT pathway is crucial for FDC differentiation (37–39). We consider that FDC becomes the main source of CXCL13 in TLSs after the FDC network is formed, similar to that in SLOs.

The CXCL13-producing PD-1 high CXCR5 negative CD4^+^ T cells we reported in RA (16, 17) do not express CXCR5, a marker typical of follicular helper T cells (Tfh). In the current study, the CD4^+^ T cells in which CXCL13 expression was induced were PD-1 high CXCR5 negative. The CXCL13-producing CD4^+^ T cells reported in breast cancer (34) had similar characteristics. The comprehensive analysis of blood and synovial samples of RA patients was used to propose a pathogenic PD-1^high^ CXCR5^-^ CD4 subset as peripheral helper T cells (Tph) (40, 41). Tph cells express factors that enable B-cell help, including IL-21, CXCL13, and ICOS. Similar to PD-1^high^-CXCR5^+^ Tfh, Tph cells induce plasma cell differentiation *in vitro* through IL-21 secretion and SLAMF5 interactions (40). In this context, CXCL13-producing CD4^+^ T cells not only promote the initial formation of TLSs but may also support anti-tumor antibody responses by B cells. Evidence for this was shown in our study, where B-cell lineages were clearly increased in patients with TLS and were associated with an improved prognosis. However, further research related to the co-localization of antigen-specific B cells with Tph cells in TLSs is warranted at the molecular level.

Proinflammatory conditions involving TGF-β promoted the differentiation of CXCL13-producing CD4^+^ T cells in our previous studies (16, 17) and in breast and ovarian cancers (33, 34). Resident fibroblasts and macrophages, and infiltrating Tregs produce TGF-β locally (42). Previously, we reported that the TGF-β signaling pathway was activated in advanced ovarian cancer and promoted tumor progression and metastasis (22). Indeed, high levels of TGF-β1 were detected in the conditioned medium of DK-09, a cell line we established from the ascites of a recurrent multidrug-resistant HGSC patient. In this study, we performed a CXCL13 induction assay using PBMCs from a healthy donor and found that TGF-β promoted CXCL13 secretion, although there was a significant difference in response to TGF-β between CD4^+^ T cells and CD8^+^ T cells. Furthermore, CXCL13 production was enhanced in CD4^+^ T cells under an IL-2 restricted environment and in CD8^+^ T cells under an IL-12 rich environment.

In addition to TGF-β, IL-2 is involved in the differentiation of CD4^+^ T cells. Quenching IL-2 by Treg or dendritic cells was reported to contribute to the differentiation of Tfh and Th17 cells (43, 44). In our previous studies, an IL-2 neutralizing antibody enhanced CXCL13 production by PD-1^high^ CXCR5^-^ CD4^+^ T cells in RA (16, 17). Gu-Trantien et al. reported that IL-2 deprivation was critical for the production of CXCL13 and the accumulation of activated Tregs in parallel with CXCL13^+^CD4^+^ TIL in breast cancer (34). However, IL-12 produced by dendritic cells and macrophages has an essential role in the interactions between the innate and adaptive immune systems. IL-12 promotes CD8^+^ cytotoxic T cell activation and expansion (45–47). Our findings that the secretory environment of CXCL13 was different between CD4^+^ and CD8^+^ T cells is interesting and consistent with the different predominance of cells expressing CXCL13 between TLSs and TILs in ovarian cancer tumor sections. The environment of CXCL13 production differs between CD4^+^ and CD8^+^ T cells, and their roles in the formation and maintenance of TLSs may be different, and should be studied in future experiments. Last, we evaluated whether TLS was induced by CXCL13. In a mouse model of spontaneously developing gastric cancer by activated STAT3 signaling, chemokines, CXCL13, CCL19, and CCL21 were induced simultaneously with tumorigenesis and TLS formation (48). We administered mouse rCXCL13 one day after tumor inoculation and succeeded in inducing TLSs. CXCL13 was highly expressed and corresponding with TLSs. CXCL13 signals B cells to enhance LT production (37, 49), and the exogenous CXCL13 may have promoted a positive feed-forward loop. In the CXCL13 treated group, the infiltration of CD8^+^ T cells around the TLS was clearly increased, and the survival time of tumor-bearing mice was also prolonged. Direct antitumor effects by CXCL13 were also observed in a colon cancer model. However, tumor growth was accelerated in CXCR5 or Rag1 knockout mice (30). The CXCL13 axis is a functional part of the relevant immune control and the TME can be altered by inducing CXCL13 and TLSs. Accordingly, our results clearly demonstrate that the induction of CXCL13 and TLSs has potential as an immune-modulatory target for ovarian cancer.

Taken together, CXCL13 is a strong prognostic factor for ovarian cancer, and is highly involved in the formation of TLS. CXCL13-producing CD4^+^ T cells induced by TGF-β under an IL-2 restricted tumor environment are important for the initial formation of TLS. The presence of TLS mobilizes various lymphocytes, and in particular, the simultaneous infiltration of B-cell lineages that are critical for the cytotoxic response of CD8^+^ T cells in ovarian cancer. The strong interaction between humoral and cellular immunity in the antitumor response was revealed, and the possibility of TLS-mediated interactions was demonstrated. *In vivo* experiments revealed the TME can be altered by inducing CXCL13 and TLSs, which might be an important immunomodulatory method to enhance antitumor immunity.

## Acknowledgement

We thank all the members of the Center for Anatomical, Pathological and Forensic Medical Research and Medical Research Support Center, Graduate School of Medicine, Kyoto University for preparing microscope slides.

## Contributors

MU designed and performed the experiments, analyzed the data, and wrote the manuscript. JH and HY designed and directed the study and edited the manuscript. ST performed statistical analyses and commented on the manuscript. HN and KA provided the clinical data. YH assisted with the experiments. AU, HS, YF, MT, KYaman, KA, KYamag, TB, NM, AY, HU, and MM provided advice on the experiments and commented on the manuscript.

## Ethics approval and consent to participate

This study was approved by Kyoto University Graduate School and Faculty of Medicine, Ethics Committee (G531), Ethics Committee of Kindai University Faculty of Medicine (27–182), and Ethics committee of the National Hospital Organization Kyoto Medical Center (19-081). Informed consent was obtained in the form of opt-out on the Web site for the patients at Kyoto University and Kindai University. Written informed consent was obtained from the patients at Kyoto Medical Center. Animal experiments were approved by the Kyoto University Animal Research Committee. This study was conducted according to Declaration of Helsinki principles.

## Date availability statement

The data analyzed in this study were previously deposited in the Gene Expression Omnibus (GEO) at GSE39204 and GSE55512 by our laboratory. All data relevant to the study are included in the article or uploaded as supplementary information.

