## Supplementary Materials for "Tertiary lymphoid structures induced by CXCL13-producing CD4^+^ T cells increase tumor infiltrating CD8^+^ T cells and B cells in ovarian cancer"

- **Supplementary Methods**
- **Supplementary Table S1-S2**
- **Supplementary Figure S1-S3**

### Supplementary Methods

#### Establishment of a patient derived cell line, DK-09

The patient was diagnosed with peritoneal carcinoma stage IIIB, and the histology indicated high grade serous carcinoma. The patient underwent primary debulking surgery and six cycles of dose-dense paclitaxel/carboplatin as adjuvant chemotherapy. The patient was treated with gemcitabine/carboplatin + bevacizumab and pegylated liposomal doxorubicin after recurrence, but they were ineffective. When cell-free and concentrated ascites reinfusion therapy (CART) was performed for the palliation of symptoms, a sample of ascites was obtained with the patient's informed consent. The cells were collected by centrifugation, lysed with ACK lysing buffer (Cat# 10-548E, LONZA, Basel, Switzerland) to remove red blood cells, and treated with Liberase (37°C for 30 minutes) (Cat# 5401127, Roche, Basel, Switzerland). CD45-MACS beads (Cat# 130-052-301, Miltenyi Biotec) were added and $CD45^{+}$ cells were removed by the column method.

Flow cytometry was used to confirm the cells were negative for CD45 after the removal of $CD45^{+}$ cells. Anti-fibroblast antibody was used to confirm the absence of fibroblasts. We confirmed that approximately 60% of the cells were positive for FOLR1, which is positive in most ovarian cancers. The cells were then passaged in RPMI + 20% fetal bovine serum, and autonomous growth was observed. In Passage 28 cells, 98.7% of the cells were CD45-negative and 77% of the cells were FOLR1-positive, and these cells were used in subsequent experiments.

**Supplementary Table S1. Patients' characteristics**

| | Kyoto Univ. cohort | Kindai Univ. cohort | Total |
|---|---|---|---|
| Histology | High grade serous carcinoma | | |
| No. of cases | 62 | 35 | 97 |
| Age (median, years) | 56.5 | 63 | 60 |
| (range) | (35-82 ) | (42-84) | (35-84) |
| Stage I | 7 | 7 | 14 |
| Stage II | 5 | 3 | 8 |
| Stage III | 44 | 17 | 61 |
| Stage IV | 6 | 8 | 14 |
| TLS positive cases | 39 | 22 | 61 |
| ( % ) | 62.9 | 62.8 | 62.9 |
| Median follow up (month) | 76 | 52 | 59 |
| (range) | (8-196) | (1-120) | (1-196) |

**Supplementary Table S2. Materials used for immunohistochemistry, RNA in situ hybridization, flow cytometry, and ELISA**

| Immunohistochemistry | Source | Catalog # |
|---|---|---|
| Mouse anti-human CD8 (Clone: C8/144B) | Nichirei Biosciences | 413201 |
| Rabbit anti-human CD4 (clone: SP35), 1:100 dilution | Cell Marque | 104R-14 |
| Mouse anti-human CD20 (clone: L26), 1:500 dilution | Novus Biologicals | NBP2-44743 |
| Rabbit anti-human CD38 (clone: SP149), 1:100 dilution | Arigo Biolaboratories | ARG52764 |
| Mouse anti-human CD21 (clone: 1F8), 1:10 dilution | Novus Biologicals | NBP1-22527 |
| Rat anti-mouse CD4 (clone: H129.19), 1:200 dilution | BD Pharmingen/BD Biosciences | 550278 |
| Rat anti-mouse CD8 (clone: YTS169.4), 1:200 dilution | Abcam | ab22378 |
| Rat anti-mouse CD19 (clone: 6D5), 1:200 dilution | Abcam | ab25232 |
| Rat anti-mouse/human Ki67 (clone: SolA15), 1:100 dilution | Thermo Fisher | 14-5698-80 |
| Rabbit anti-mouse CD8 (clone: EPR20305), 1:2000 dilution | Abcam | ab209775 |

| RNA in situ hybridization | Source | Catalog # |
|---|---|---|
| Hs-CXCL13 (for human) (C1) | Advanced Cell Diagnostics | 311321 |
| Mm-Cxcl13 (for mouse) | Advanced Cell Diagnostics | 406311 |
| Hs-CD4 (C2) (for human) | Advanced Cell Diagnostics | 605601-C2 |
| Hs-CD8A (C3) (for human) | Advanced Cell Diagnostics | 560391-C3 |
| Hs-CR2 *(C2) (for human) | Advanced Cell Diagnostics | 859071-C2 |
| Opal 690, 1:1000 dilution | PerkinElmer | FP1497001KT |
| Opal 570, 1:1500 dilution | PerkinElmer | FP1488001KT |

*Hs-CR2 targeting 269-1303 of NM_001006658.3

| Flow cytometry | Source | Catalog # |
|---|---|---|
| APC-conjugated mouse anti-human CXCL13 (clone: 53610) | R&D Systems | IC801A |
| BV421-conjugated rat anti-Human CXCR5 (clone: RF8B3) | BD Biosciences | 562747 |
| PE-conjugated mouse anti-Human CD279 (clone: EH12.2H7) | BioLegend | 329906 |

| ELISA | Source | Catalog # |
|---|---|---|
| Human CXCL13/BLC/BCA-1 Quantikine ELISA Kit | R&D Systems | DCX130 |
| TGFβ-1 Human ELISA Kit | Thermo Fisher | BMS249-4 |

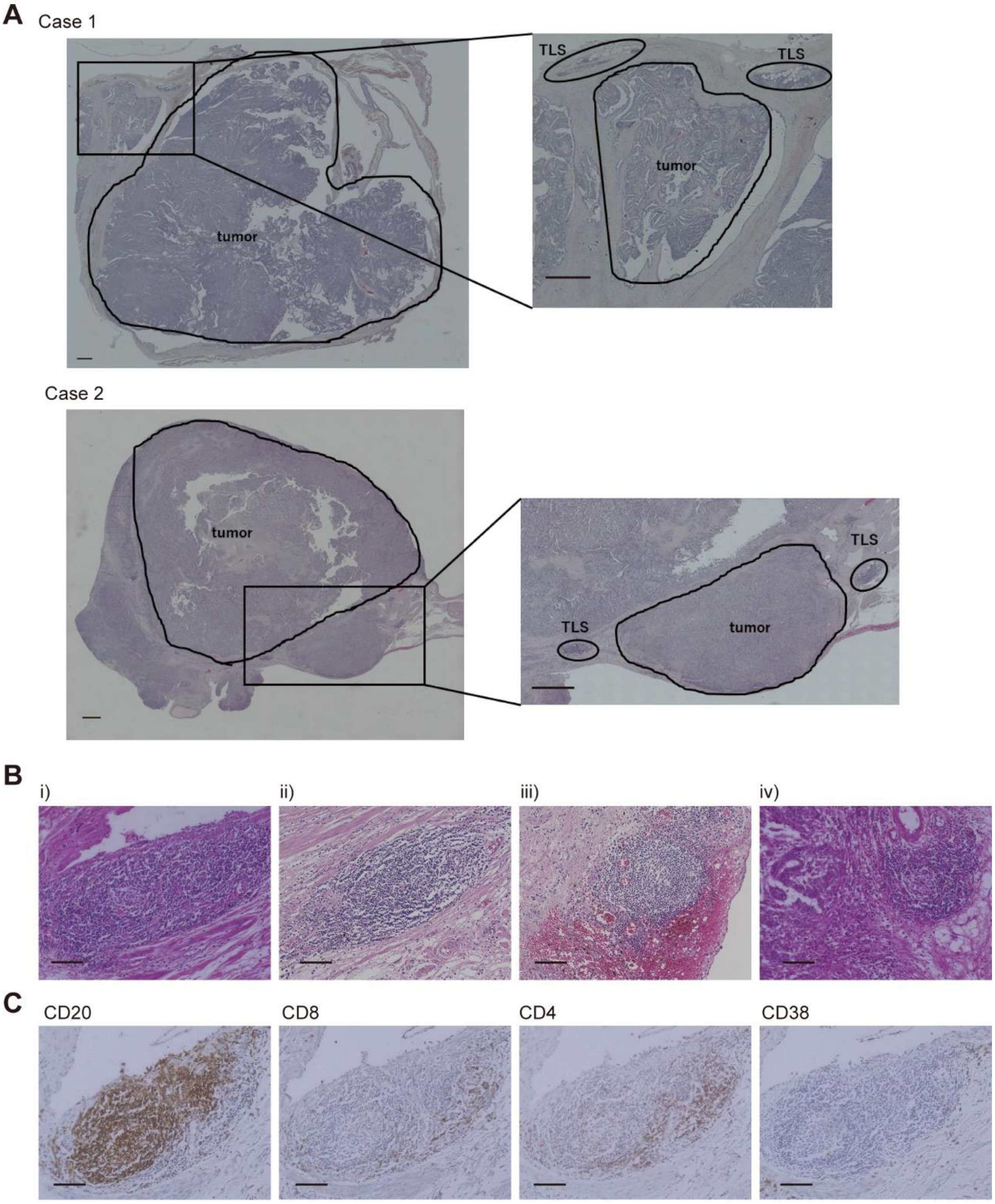

**Supplementary Figure S1. Representative TLS and dissection of TLS and tumor sites (TIL) for repertoire analysis.**

**(A)** As shown in the figure, the tumor area and TLS area were separated by microdissection. The tumor region was analyzed as TIL and the TLS region as TLS for repertoire analysis. **(B)** Representative H&E images of TLS. **(C)** Distribution of various lymphocytes in TLS. IHC images (DAB) of CD20, CD8, CD4, and CD38 in A(i). Scale bars indicate 1000 μm in (A), and 100 μm in (B) and (C).

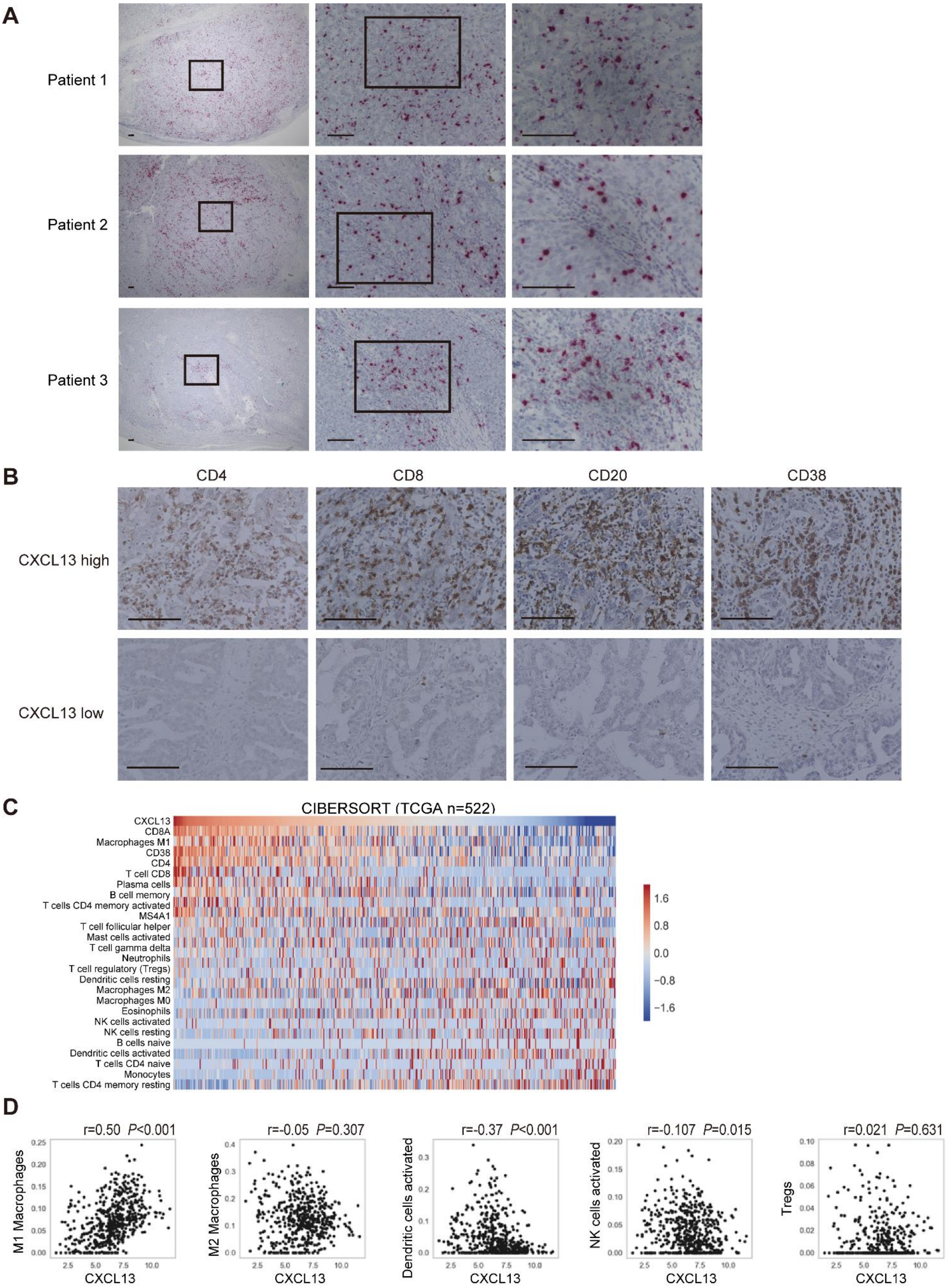

**Supplementary Figure S2. CXCL13 expression and intratumor infiltrating lymphocytes.**

**(A)** CXCL13 expression in tumor infiltrating immune cells by RNA ISH (Fast RED). Representative three cases with a high expression of CXCL13 in immune cells. Scale bars indicate 100 µm. **(B)** CXCL13 gene expression and distribution of several types of tumor-infiltrating immune cells. The upper row shows a case with high CXCL13 gene expression and the lower row shows a case with low CXCL13 gene expression. Scale bars indicate 100 µm. **(C)** The distribution of infiltrating immune cells into the tumor site and CXCL13 gene expression using CIBERTSORT (n=522). Overall view of the image shown in Figure 2D. **(D)** Correlation of CXCL13 gene expression with M1- or M2-macrophages, activated dendritic cells, activated natural killer cells, or regulatory T cells (Tregs) in CIBERSORT. Correlation was determined by Spearman's correlation test.

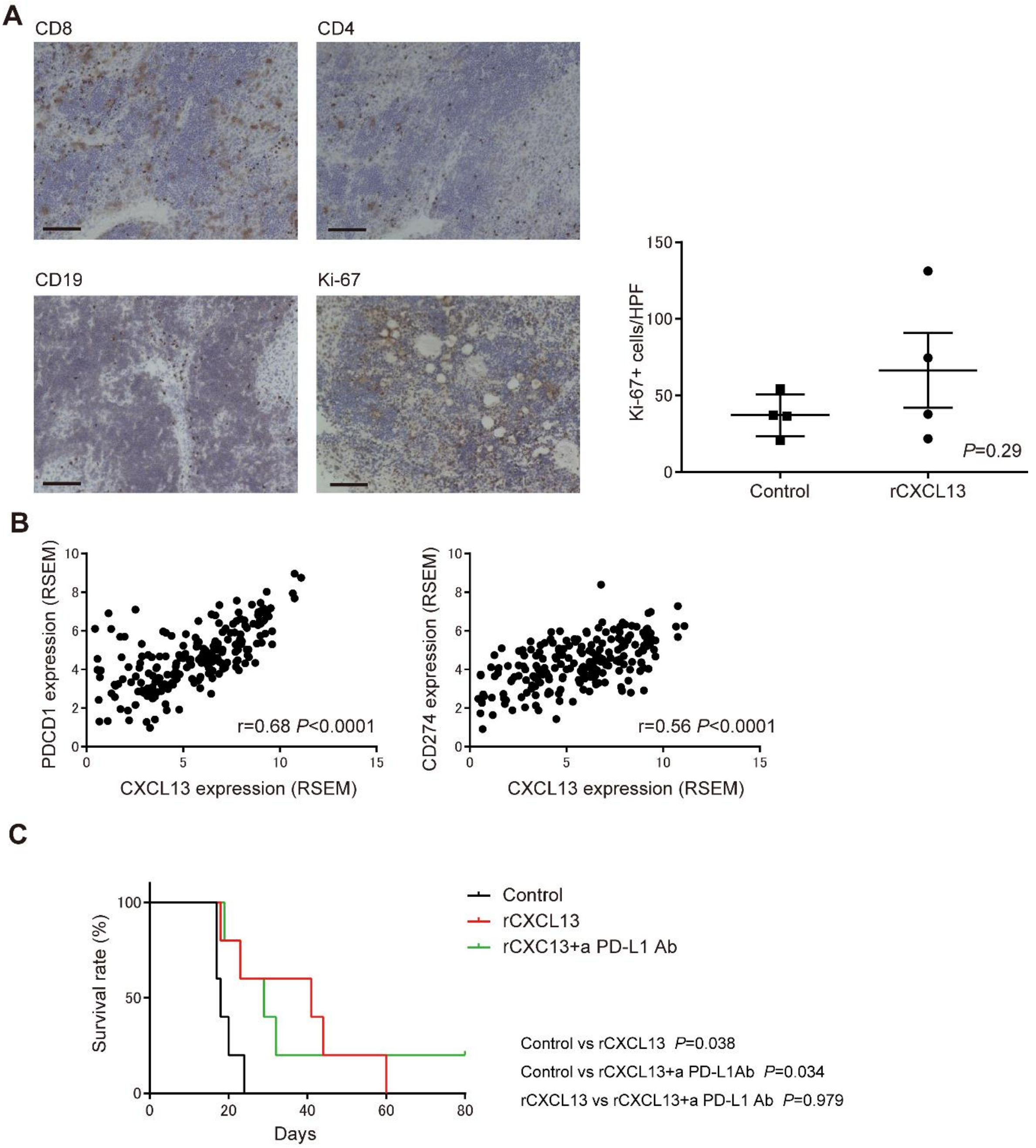

**Supplementary Figure S3. Anti-tumor immunological effects of CXCL13 in a mouse ovarian cancer model**

**(A)** Analysis of cells in the TLS induced by rCXCL13 in mice by IHC of CD8, CD4, CD19, and Ki-67 (DAB). Scale bars indicate 100 μm. The number of Ki-67 positive cells (mean count of five sections at high power field) in TLS was compared between the rCXCL13 treated group and the control group. Data are shown as the mean ± SEM of four samples. Statistical significance was determined by two-tailed Student's *t*-test. **(B)** Correlation between CXCL13 and PD-1 (PDCD1) or PD-L1 (CD274) expression in TCGA (n=217). Correlation was determined by Spearman's correlation test. **(C)** Survival curves of tumor-bearing mice treated with rCXCL13 or rCXCL13 + anti-PD-L1 antibodies. Analyses were performed using Kaplan-Meier estimates and log-rank tests. a PD-L1 Ab indicates anti PD-L1 antibody in (C).
